# Clinical progression of clonal hematopoiesis is determined by a combination of mutation timing, fitness, and clonal structure

**DOI:** 10.1101/2025.02.28.640879

**Authors:** Eric Latorre-Crespo, Neil A. Robertson, E. Gozde Kosebent, Louise MacGillivray, Lee Murphy, Mesbah Uddin, Eric Whitsel, Michael Honigberg, Alex Bick, Alexander P. Reiner, Valeria Orrù, Michele Marongiu, Francesco Cucca, Edoardo Fiorillo, Ian J. Deary, Sarah Harris, Simon Cox, Riccardo Marioni, Linus Schumacher, Tamir Chandra, Kristina Kirschner

**Author notes:** Equal Contribution (Eric Latorre-Crespo, Neil A. Robertson).

## Abstract

Clonal hematopoiesis (CH) is characterized by expanding blood cell clones carrying somatic mutations in healthy aged individuals and is associated with various age-related diseases and all-cause mortality. While CH mutations affect diverse genes associated with myeloid malignancies, their mechanisms of expansion and disease associations remain poorly understood. We investigate the relationship between clonal fitness and clinical outcomes by integrating data from three longitudinal aging cohorts (n=713, observations=2,341). We demonstrate pathway-specific fitness advantage and clonal composition influence clonal dynamics. Further, the timing of mutation acquisition is necessary to determine the extent of clonal expansion reached during the host individual’s lifetime. We introduce MACS120, a metric combining mutation context, timing, and variant fitness to predict future clonal growth, outperforming traditional variant allele frequency measurements in predicting clinical outcomes. Our unified analytical framework enables standardized clonal dynamics inference across cohorts, advancing our ability to predict and potentially intervene in CH-related pathologies.

## Introduction

We are born with an estimated 50,000 to 200,000 hematopoietic stem and progenitor cells (HSPCs) that are responsible for constituting the blood system across our lifespan – a remarkably small number given the high rate of output of differentiated blood cells maintained over this time ^1^. From the perspective of somatic evolution, this demarks the origin and period over which mutation selection can perturb the hematopoietic system.

The growth rate – or fitness – of HSPC clones is defined as the proliferative advantage over cells carrying no or only neutral mutations ^2^. The study of CH in large cross-sectional cohorts has provided a wealth of perspectives on the genetic drivers, their prevalence, and associations with numerous clinical features ^3–5^. However, cross-sectional studies leave a multitude of questions regarding how CH develops, the dynamics of clonal growth, and how clonal fitness might interact with cell-extrinsic determinants as we age. In 2022, we and others generated the first large-scale longitudinal studies of CH which quantified trajectories of mutations across a range of genes, highlighting similar dynamics across several independent cohorts ^6–8^. We identified gene-specific fitness effects across the 7th to 9th decades of life which, for some genes, can outweigh inter-individual variation ^6^.

Here, we have annotated CH status across three longitudinal cohorts in 713 participants aged 55-99, totaling 2,341 observations. We find that mutations in the same gene with different clonal structures grow with different fitness and that there is a strong correlation between inferred fitness and the estimated time of mutation acquisition. Moreover, different clones competing for space in an individual’s hematopoietic compartment led to alterations in clone size distribution that affect the fitness inference and their predicted future growth. These findings highlight the importance of incorporating personalized clonal dynamics in the development of clinical risk predictors for CH.

## Results

### Three longitudinal cohorts of aging exhibit concordant clonal hematopoiesis status

Previous reports have linked clone size measured in variant allele frequency (VAF) to outcomes such as all-cause mortality, leukemia risk, and cardiovascular disease ^4,5^. Recently, we and others have associated the most common CH genes with distinct fitness effects ^6–8^. However, the associations of these fitness effects with clinical outcomes remain unknown. To generate a data set of sufficient size to elucidate the relationship between clonal fitness and clinical outcomes we used error-corrected target sequencing as previously described on an additional 63 Lothian Birth Cohort (LBC) participants. We then combined this data with published data from three longitudinal cohorts of aging (SardiNIA, Lothian Birth Cohorts of 1921 and 1936 and Women’s Health Initiative (WHI)) (Fig. 1A, Supplemental Table 1) ^6–8^.

**Figure 1:**
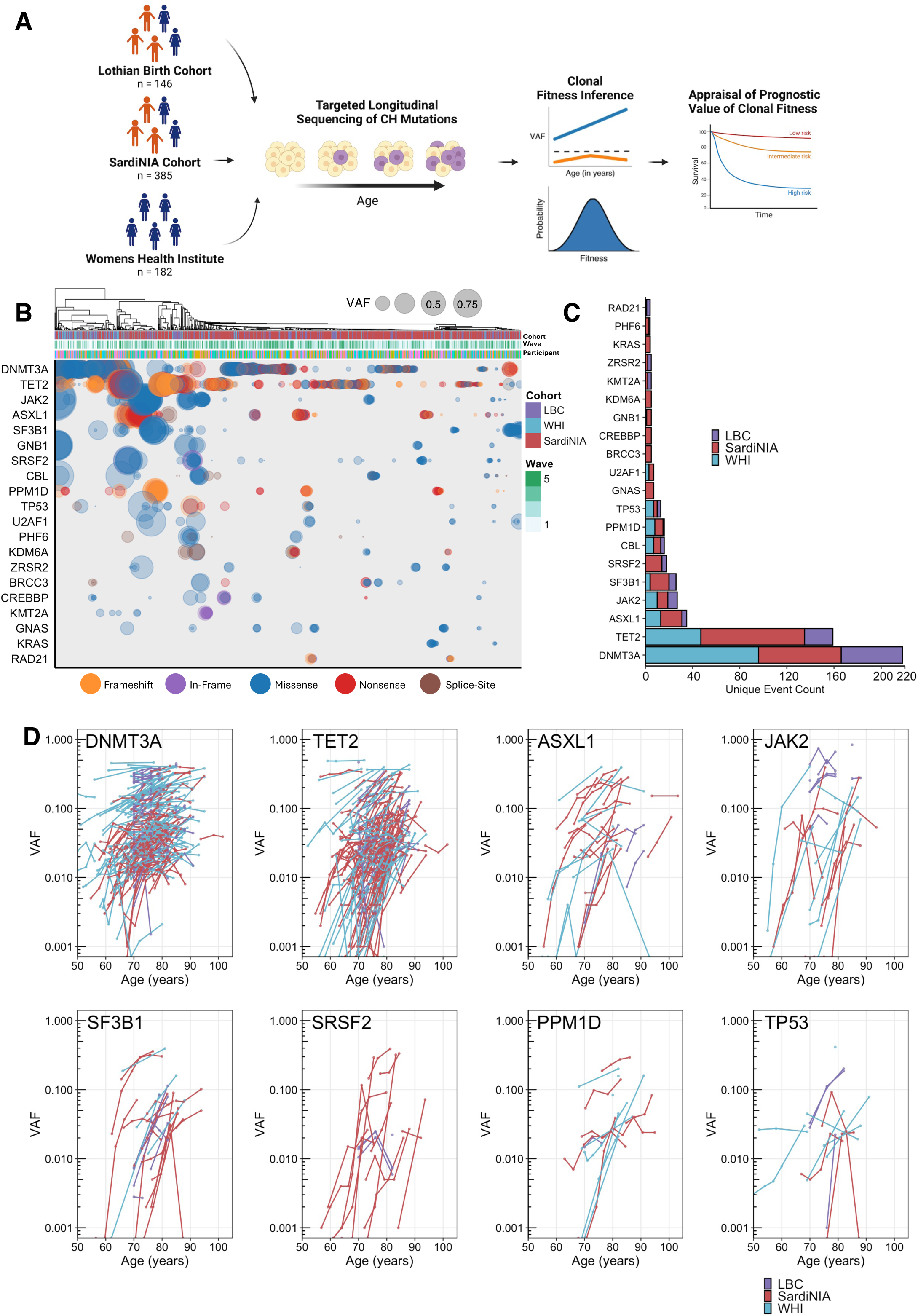
Three longitudinal cohorts of aged individuals exhibit concordant clonal hematopoiesis status. **A.** Schematic showing the targeted sequencing workflow for three longitudinal cohorts - the Lothian Birth Cohort (LBC), the Sardinian Longitudinal Project (SardiNIA), and the Women’s Health Institute (WHI); with n, the number of participants of each cohort. **B.** Variant allele fractions in the 20 most frequently mutated genes, showing samples from all participants across all time points clustered by the largest VAF of a mutation within a given gene. Each dot indicates the presence of the largest mutation within a gene for a given individual, scaled by VAF and colored by predicted functional consequence. The annotation bar indicates the participant, cohort, and wave number of samples. **C.** Counts of mutations in each of the top 20 genes, with bar stacks colored by cohort showing the distribution across different populations. **D.** Longitudinal trajectories of variant allele frequencies for commonly mutated variants, colored by cohort with log10 scaled y-axis, demonstrating consistent patterns of clonal expansion across cohorts.

The Lothian Birth Cohorts of 1921 and 1936 (LBC1921 and LBC1936, respectively) are two prospective studies designed to assess the physiological determinants of cognitive aging and were recruited from the Lothian region of Scotland ^9,10^. The Sardinian Longitudinal Project (SardiNIA) aimed to understand factors influencing lifespan using a healthy, long-lived, and genetically homogenous founder population ^7,11^. Finally, the Women’s Health Institute (WHI) cohort was devised to capture the long-term health effects of aging in post-menopausal women ^12^. The combination of the three cohorts studied herein represents populations with significantly divergent age and sex distributions, and geographical and genotypic backgrounds (age range 55-99 years at baseline with a total number of 713 participants across all cohorts (Fig. S1A, S1B)). Despite this heterogeneity, we were able to recapitulate the canonical enrichment patterns of CH somatic driver events across all cohorts (Fig. 1B-C).

To improve the quality of the data sets and account for sequencing noise, mutation trajectories in the LBC were filtered using our previously described LiFT method ^6^. LiFT was not applicable to the other two cohorts as unfiltered sequencing data were unavailable. This was followed by manual curation across all cohorts to reduce over-enriched sequencing artifacts or those that display anomalous VAF levels across the time series (see Supplemental Methods), resulting in a majority of trajectories that were increasing in VAF over time (Fig. 1D) ^6^.

In concordance with previous reports, we found mutations in DNMT3A, TET2 and ASXL1 being the most frequent mutations across all three cohorts (Fig. 1B, 1C, S1E, S1F) with VAF across all somatic variants showing a moderate increase with age (R^2^= 0.003, p=1.055 e-01) (Fig. 1D, S1D, S1G) ^4,5^. The most commonly found type of mutations were missense mutations, followed by frameshift and nonsense mutations across all cohorts (Fig. S1C). These findings demonstrate that despite differences in cohort composition, geographical location, and data acquisition, mutational CH status and prevalence are robust measurements and independent of cohort specifics. Overall, we harmonized three diverse longitudinal cohorts to generate a coherent dataset to interrogate CH status and its clinical associations.

### A cross-cohort unified analysis recapitulates gene-specific fitness effects

We developed a computational framework that leverages longitudinal VAF trajectories to determine fitness, clonal structure, and ‘Age at the Time of Mutation Acquisition’ (ATMA) across all cohorts (Fig. 2A, Methods). Briefly, our pipeline uses a hidden Markov model to determine the posterior fitness distributions for all mutations per participant and possible clonal structures. The most likely clonal structures and fitness estimates are then used to estimate the ATMA, enabling predictions of historical and future clonal dynamics.

**Figure 2:**
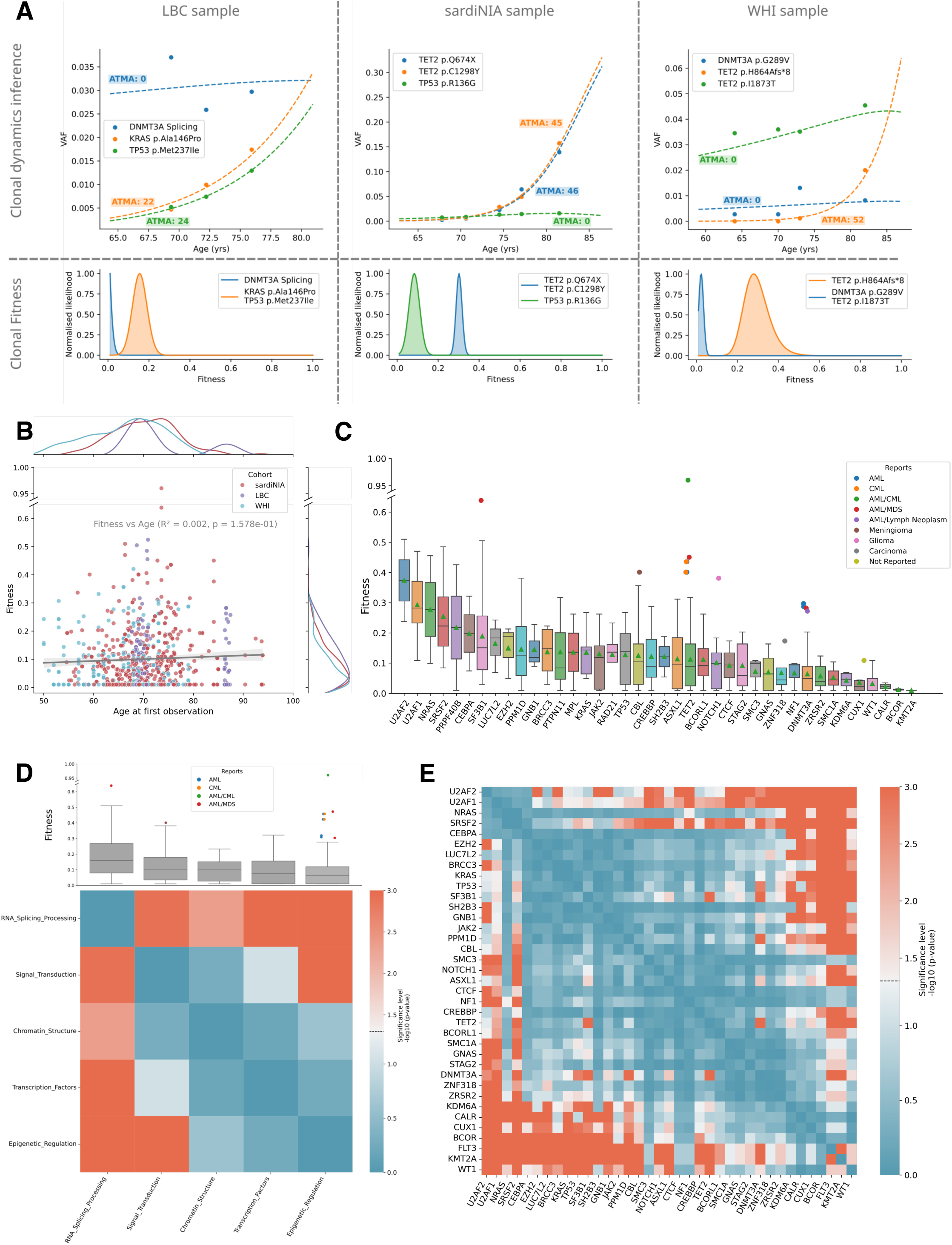
A cross-cohort unified analysis recapitulates gene-specific fitness effects. **A.** Mutation trajectories (dots) and deterministic model fits (dashed lines) for three example participants, one from each cohort (top row). Growth curves shown are deterministic fits using the maximum of the posterior distribution of fitness (bottom row). Posterior distributions are shown per clone, and the legend indicates which mutations, according to the inference, are likely co-occurring on a clone. ATMA: Age at Time of Mutation Acquisition. **B.** Clone fitness versus participant age at first observation, with points colored by cohort. Marginal distributions shown as normalized Kernel Density Estimation (KDE) curves on each axis. **C.** Fitness of mutations in all three cohorts grouped by gene they occur in, ranked by mean fitness (green triangles). Boxes show median and exclusive interquartile range. Outlier points are colored according to previously reported associations with malignancies (see legend). **D.** Fitness distribution across gene functional categories (Epigenetic Regulation, RNA Splicing, Signal Transduction, Transcription Factors, and Chromatin Structure, see Methods). Boxes indicate exclusive interquartile range with median and points show outliers. Below, heatmap displays statistical significance (-log10 p-values) of pairwise Brunner-Munzel tests between categories, with red colors indicating stronger significance. **E.** Pairwise comparison of fitness across genes. Heatmap displays statistical significance (-log10 p-values) of pairwise Brunner-Munzel tests between categories, with red colors indicating stronger significance.

Our fitness inferences across cohorts revealed several key insights. First, we found no correlation between fitness and participant age at initial observation (R²=0.02, p=0.16, Fig. 2B). While we observed minor cohort-specific differences in fitness effects, most notably between LBC and WHI (Brunner Munzel test, BM=-3, p=2.384e-03) and between SardiNIA and WHI (BM=-2, p=2.948e-02, Fig. S2A), these differences disappeared when comparing only the maximum fitness observed per participant (Fig. S2B). This suggests that cross-cohort variations in fitness distribution primarily reflect differences in detection sensitivity for low-fitness variants that remain at low VAF, supporting the robustness of our fitness predictions across cohorts.

Consistent with our previous report, we observed significant gene-specific variations in fitness distributions (Fig. 2C, S2C) ^6^. Overall, splicing gene mutations showed the highest fitness advantage (mean of 0.20), with U2AF2 and U2AF1 displaying the highest mean fitness estimates of 0.37 and 0.29, respectively (Fig. 2C-E; see Methods for gene categories). In contrast, epigenetic regulators showed the lowest fitness advantage (mean of 0.09), with DNMT3A and TET2 displaying fitness estimates of 0.06 and 0.11, respectively (Fig. 2C-E). Encouragingly, high-fitness variants within the epigenetic regulators category corresponded to previously identified cancer driver mutations, supporting our ability to detect likely pathogenic variants at early stages of growth (Fig. 2C,). We found that among all early-stage variants (VAF<10%) with high fitness scores (>0.3), 46% (12/26) were previously documented in leukemia (Fig. S2D).

### Clonal composition drives differential fitness effects

To further understand the connections between fitness and mutated genes, we investigated how the hierarchical arrangement of somatic mutations within HSPCs, or clonal structure, influences the longitudinal evolution of variants. Clonal structure - encompassing both their internal mutational composition and the simultaneous growth of co-occurring variants within the blood has emerged as a critical determinant of clonal dynamics ^13,14^. To quantify the effect of mutations occurring in the same individual but on separate clones, we modeled clonal evolution in each individual across the cohorts based on the inferred fitness and ATMA. Mutational context within an individual, i.e., which other mutations are present and whether these are on separate clones or co-occurring, needs to be taken into account for accurate fitness predictions as demonstrated in the following example: We identified two competing clones in the same individual, with one clone carrying a *DNMT3A T735C* mutation and another clone likely harboring both a *DNMT3A A326G* and a *JAK2 V617F* mutation (Fig. 3A). While isolated modeling predicted all mutations would reach VAF 0.5 by the maximum human life span (age 120), observed trajectories revealed suppressed proportional growth, i.e., growth in VAF, of the clone carrying two mutations, highlighting the importance of accounting for *in vivo* clonal competition (Fig. 3A). Across all individuals, this comparative analysis showed a difference in predicted mean VAF at age 120 of 10% between the isolated growth and mutational context models, with 13% of participants exhibiting differences in VAF predictions exceeding 20% (Fig. 3A, B). These findings demonstrate that accurate prediction of variant evolution requires consideration of the complete clonal composition within an individual, as competition for space in the hematopoietic compartment among coexisting clones substantially influences their trajectories over their lifespan.

**Figure 3:**
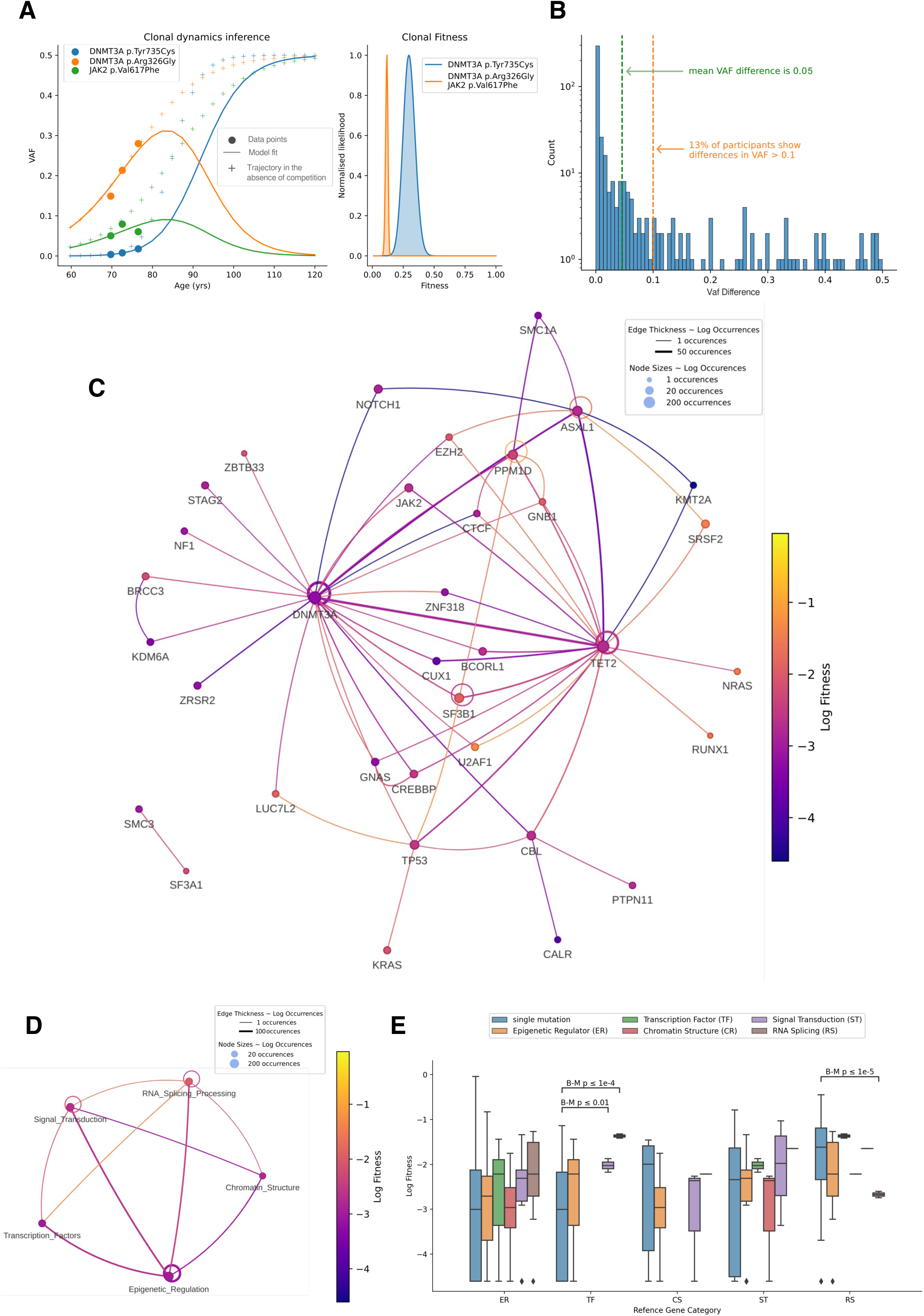
Clonal composition drives differential fitness effects. **A.** Example clonal expansion trajectories demonstrating the impact of clonal competition in a participant with two independent clones - one with DNMT3A T735C mutation (blue) and another likely harboring both DNMT3A A326G (orange) and JAK2 V617F mutations (green). (Left) Points show observed VAF, solid lines show deterministic evolution predicted with a model accounting for clonal composition (context-aware model), and crosses show predictions from an isolated growth model. The difference between solid lines and crosses illustrates the importance of mutational context in predicting the change of trajectories. (Right) Posterior distributions are shown per clone and the legend indicates which mutations, according to the inference, are likely co-occurring on a clone. **B.** Distribution of differences in predicted VAF at age 120 between the isolated and context-aware models across all participants and mutations. **C.** Network visualization showing mutation co-occurrence patterns. Nodes represent individual mutations, with node size proportional to log counts of mutation instances and node color indicating average mutation log fitness (scaled between minimum and maximum log fitness values). Edges connect co-occurring mutations in a clonal structure, with edge width proportional to co-occurrence log counts and edge color representing the average log fitness of connected mutations. **D.** Same as in **C.** but with mutations classified into gene categories. **E.** Fitness of mutations across different combinations of gene categories. For each category (ref) we show boxplots comparing the log fitness of singly occurring mutations in a clone and co-occurring with mutations in any category. Boxes are colored by category and show median and exclusive interquartile range. Statistical significance of fitness differences between isolated and co-occurring mutations was assessed for log fitness using the Brunner-Munzel test with Benjamini-Yekutieli correction for multiple comparisons.

To address variant cooperation within a clone, we examined differences in fitness between mutations within a gene depending on co-occurring mutations in our inferred clonal compositions. We found that clones with co-occurring mutations have different fitness than clones with isolated mutations in the same genes (Fig. 3C, S3A-G). While such context-dependent fitness differences were common across most genes, statistical power was limited for genes other than epigenetic regulators due to the rarity of co-mutations. For ASXL1, we observed that the combination of ASXL1/SRSF2 mutations in the same clone conferred greater fitness advantages than ASXL1/EZH2 (0.27 and 0.19 respectively with both n=1) with the lowest observed mean log fitness when ASXL1 mutations co-occur with TET2 mutations (mean fitness 0.06, n=27) (Fig. S3D and S3G). These findings are consistent with the known association of prognosis in acute myeloid leukemia (AML) patients with co-mutated ASXL1/SRSF2 and ASXL1/EZH2 clones ^15,16^. In contrast, PPM1D’s highest fitness effects were observed in a clone harboring multiple PPM1D mutations (fitness 0.36, n=1), followed by co-occurrence with GNB1 (fitness 0.24, n=1) and SF3B1 (fitness 0.19, n=1) (Fig. S3C and S3G). These findings are particularly relevant given PPM1D’s role in therapy-related secondary leukemias, especially following treatment for solid cancers ^17,18^.

We also analyzed how mutations in different gene categories interacted by comparing the fitness effects of single mutations versus co-occurring mutations across categories (Fig. D-E, see Methods). While co-occurring mutations were relatively rare, we found that single mutations in RNA splicing genes (n=39) conferred significantly higher fitness than when these mutations co-occurred with epigenetic regulator mutations (n=11, p<1e-5). Interestingly, though not statistically significant due to small sample size, clones with co-mutations in RNA splicing and transcription factor genes showed the highest average fitness (0.25, n=2). These findings demonstrate that fitness effects within a clone can not only be assigned on a gene level but also when considering gene categories, making clinical utility more likely.

Overall, we demonstrated that clonal composition within an individual and mutation profiles within the same clone play a major role in driving fitness effects.

### Mutation timing complements fitness in determining the clinical progression of clonal hematopoiesis

Our improved analysis of clonal evolution and our model’s ability to predict clone trajectories in an individual context enabled us to develop more precise metrics for quantifying the potential impact of mutations. Analysis of mutation fitness relative to the ATMA revealed a significant positive correlation (R=0.89, p=5.7483-281, Fig. 4A), suggesting that higher-fitness mutations accumulate with increasing age. This pattern may indicate a progressive decline in DNA surveillance mechanisms across the lifespan, allowing more deleterious mutations to accumulate in older individuals. Notably, we observed a complete absence of variants with low fitness and large ATMA which is explained by insufficient time for recently mutated clones to reach detectable levels in older participants. This observation is further supported by variants nearest to this detection threshold corresponding to the oldest study participants (Fig. S4A). While the limitation in detecting low fitness high ATMA variants may artificially inflate the observed correlation between these two parameters, the fundamental relationship demonstrating enhanced maximum fitness with increasing ATMA remains robust.

**Figure 4:**
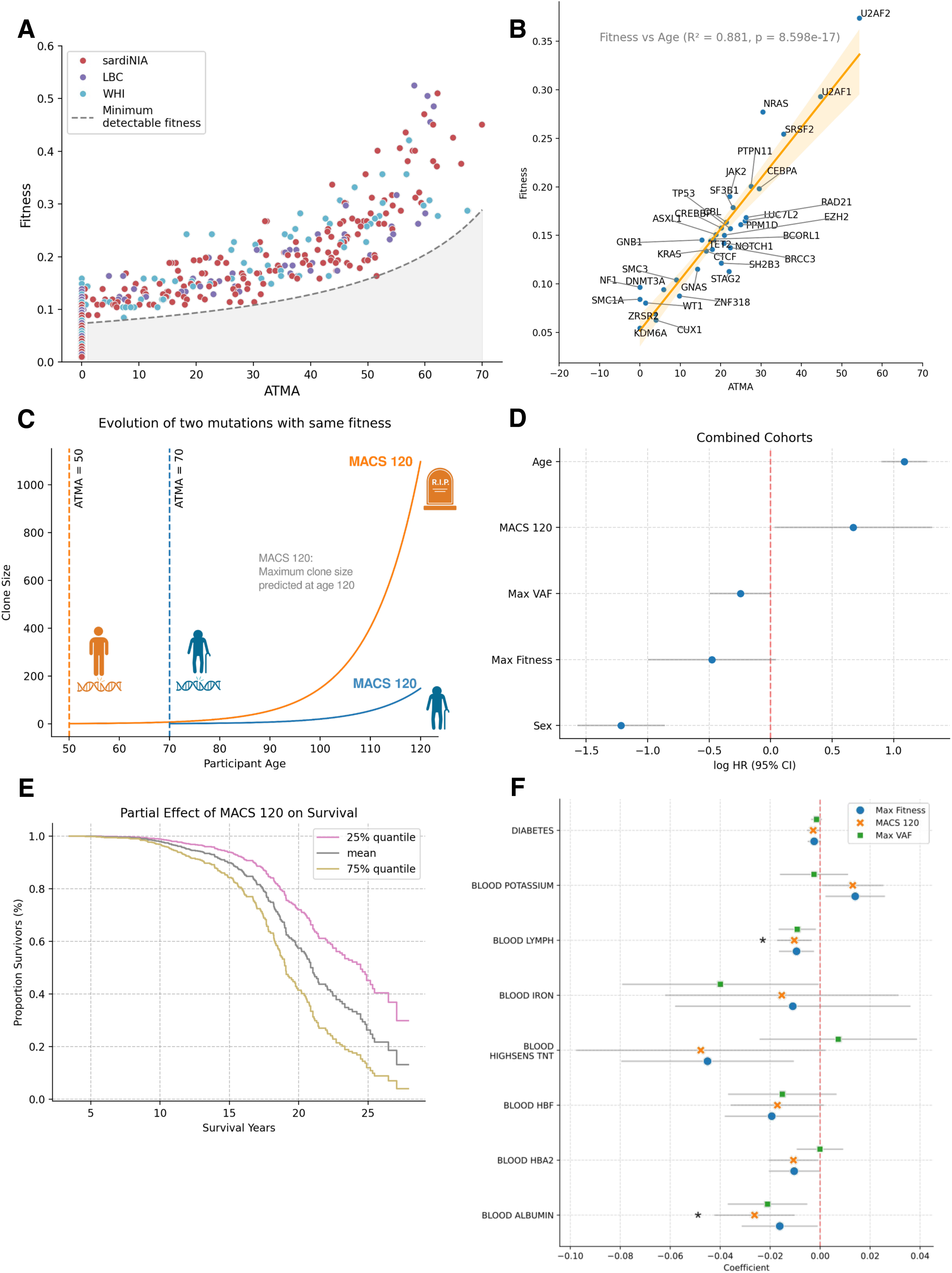
**Mutation timing complements fitness in determining clinical progression of clonal hematopoiesis.** A. Mutation fitness and Age at Time of Mutation Acquisition (ATMA) shows positive correlation. Points colored by cohort. The absence of low-fitness high-ATMA variants is due to detection limitations (see Methods). B. Gene-level analysis showing mean fitness versus mean ATMA for each gene. Orange line shows the optimal linear regression fit (R^2^=0.88, p=8.6e-17) and the shaded area confidence intervals (obtained through bootstrapping data points). C. Theoretical comparison illustrating the importance of mutation timing. Two mutations with identical fitness (s=0.1) but different acquisition times (age 50 vs 70) show dramatically different growth trajectories of clone size during the lifespan of an individual. D. MACS120 impact on survival. Cox proportional hazard ratios with 95% confidence intervals for predictive metrics: MACS120 (z-score normalized), Maximum VAF (z-score normalized), Maximum fitness (z-score normalized), Age at first observation (z-score normalized), and Sex. MACS120 shows statistically significant association with survival (PH=0.66, p=0.04). E. Partial effects of MACS120 on survival probability over time, showing how variations in MACS120 (25% quantile, baseline and 75% quantile) influence survival trajectories while controlling for other variables. F. Association between Maximum Fitness (blue dot), Maximum VAF (green square) and MACS120 (orange cross) and longitudinal changes in blood markers at the individual level. For each blood marker where any of the three metrics showed a significant correlation (see Methods) we display effect size and confidence intervals with a grey line. Acronyms: Intracellular Adhesion Molecule (ICAM), Lymphocytes (LYMPH) High-Sensitivity Cardiac Troponin T (HIGHSENS TNT), Fetal Hemoglobin (HBF), Hemoglobin Alpha 2 (HBA2).

A strong correlation between fitness and ATMA is also observable at the gene-level (R^2^=0.88, p=8.598e-17, Fig. 4B). Notably, splicing factor mutations in U2AF1 and U2AF2 showed the highest mean values for both metrics (respectively, fitness=0.37, 0.29, and ATMA=54.3, 44.7), while DNMT3A mutations display the lowest values (fitness=0.06 and ATMA=3.71). Both the early acquisition and low fitness of DNMT3A mutations and the high fitness and late acquisition of mutations affecting spliceosome genes aligns with previous observations, supporting our ability to predict ATMA ^7, 19^.

We further explored how the timing of mutation acquisition proves crucial for predicting clonal growth outcomes through a theoretical comparison of two mutations with identical fitness but different acquisition times (Fig. 4C). A mutation acquired at age 50 has sufficient time to reach potentially dangerous clone sizes within a typical human lifespan. In contrast, the same mutation acquired at age 70 never achieves comparable expansion (Fig. 4C). This observation led us to develop Maximum Clone Size at age 120 (MACS 120), a metric that predicts the theoretical maximum size any variant would reach by age 120 in an individual, which considers the effects of both growth rate and mutation timing. Unlike maximum fitness, MACS 120 shows no correlation with participant age (R²=0.09, p=0.04 and R²=0, p=0.8, respectively; Fig. S4C), suggesting it better captures intrinsic clone characteristics.

Cox proportional hazards analysis of survival data across combined cohorts revealed that while age and sex remained the strongest predictors of survival, MACS 120 demonstrated significant predictive power (log proportional hazards: LPH=0.66, p=0.04; Fig. 4D, E). In contrast, maximum VAF and maximum fitness showed no statistically significant correlation (LPH =-0.24, p=0.06 and LPH=-0.48, p=0.07, respectively, Fig. 4D, S4D-G), though this may be limited by sample size. Cohort-stratified survival analysis yielded similar results but only showed significant levels for MACS120 for the SardiNIA cohort, which was expected given the small sizes and distinct age compositions of the LBC and WHI cohorts (Fig. S4H). Further, age-stratified survival analysis showed MACS 120’s strongest association in the 70-79 age group (Fig. S4I).

To investigate clinical effects of CH beyond mortality outcomes, we analyzed associations between participant features and longitudinal rates of change in blood markers (n=43) using Linear Mixed Models (Fig. 4F). To be able to compare rates of change in markers across cohorts, we transformed and normalized marker values in each cohort to closely match a standard normal distribution (see Materials and Methods). While this procedure loses age-effects associated to the distinct ranges of age in each cohort, it allows for direct comparison of the observed longitudinal differences across cohorts. Despite the low number of available longitudinal samples for each marker (224 participants on average), we found 15 significant associations prior to multiple test correction, with 2 of them remaining significant after correction (2-stage Benjamin-Hochberg). While Max Fitness and Max VAF showed some overlapping associations, particularly with lymphocyte and albumin levels, they also demonstrated differential correlation patterns. Most notably, Max Fitness exhibited a significant positive association with potassium levels (β = 0.014, 95% CI [0.002, 0.02]), while Max VAF showed a distinct pattern in iron levels in blood (β = −0.039, 95% CI [-0.078, 0]). MACS 120 showed largely concordant associations with Max Fitness, while capturing some of the increased predictive power of Max VAF in several associations. This was showcased in the association of MACS 120 with the rate of change in lymphocyte and albumin levels which remained the only significant associations after multiple testing correction (respectively, p=0.05, β= −0.01 and p=0.05, β=-0.026). When analyzed separately for each cohort, MACS 120 was again the only metric to show significant associations with the rate of change of blood markers after multiple testing correction and displayed a further association with the rate of change of neutrophil levels in the SardiNIA cohort (p=0.05, β=0.012). This suggests that MACS 120 captures aspects of the impact of clone size and fitness on physiology not ordinarily captured by Max VAF or Max Fitness alone, providing a more comprehensive measure of CH’s systemic effects.

All three metrics - Max Fitness, MACS 120, and Max VAF - demonstrated negative associations with the rate of change of lymphocyte counts, indicating a potential impact on adaptive immunity that scales with clone size and fitness - although our analysis cannot determine the direction of causality ^20^ (Fig. 4F). Furthermore, all metrics showed significant positive correlations with the rate of change of neutrophil counts in the SardiNIA cohort, suggesting a relationship between clone fitness and myeloid expansion. Markers of organ function, on the other hand, showed varied associations across metrics. The rate of change in blood albumin levels was negatively associated with all metrics, demonstrating a broader impact on inflammatory states (Fig. 4F). Both Max Fitness and MACS 120 showed negative associations with the rate of change of diabetes markers, pointing to potential metabolic implications of CH. While several of these associations were recapitulated when analyzing each cohort separately (Fig. S4H), others emerged only when analyzing the cohorts together, highlighting the value of our unified cross-cohort approach.

These findings demonstrate that CH influences multiple physiological systems, with distinct patterns of association depending on whether clone size, fitness, or predicted growth potential is considered. The broad range of affected parameters, from immune function to metabolism and kidney function, underscores the systemic nature of CH’s impact on health and supports the utility of incorporating multiple metrics into clinical assessments.

## Discussion

The clinical significance of clonal hematopoiesis has been established through numerous cross-sectional studies demonstrating associations with adverse health outcomes ^4,5,21^. However, the field has struggled to translate these population-level associations into actionable clinical insights for individual patients. Recently, three independent research groups attempted to address these limitations through longitudinal analyses and found that these constraints may be due to a reliance on simplified metrics like VAF, which fail to capture the complex biological processes driving clonal evolution ^6–8^. Their disparate methodological approaches, however, impaired the comparison of findings and highlighted the need for a unified analytical framework. Our study represents the first comprehensive integration of these longitudinal cohorts under a single analytical approach, significantly boosting statistical power and enabling robust predictions of clinical outcomes. This unified analysis reveals that understanding clonal dynamics requires consideration of both the intrinsic properties of mutations and their co-mutational and temporal context.

The strong correlation we have observed between mutation fitness and mutation acquisition timing reveals a fundamental aspect of clonal evolution that has not been made explicit in previous studies. Early-acquired mutations, particularly in epigenetic regulators like DNMT3A, often display modest fitness effects but achieve substantial clone sizes through prolonged expansion, per previous findings ^7^. In contrast, late-acquired mutations, such as those affecting splicing factors, typically show higher fitness effects - possibly reflecting age-related deterioration in cellular quality control mechanisms. This temporal pattern helps explain the paradoxical observation that DNMT3A mutations are the most common mutations observed in CH despite their relatively low fitness effects while rarer mutations in genes like U2AF1 are more frequently associated with progression to malignancy. Of course, this interpretation is limited by the current assumption of constant fitness over past growth. Furthermore, by incorporating competitive dynamics between coexisting clones in our model, we can explain a deceleration in the growth in VAF without making assumptions about intrinsic dynamical changes in fitness. Whether fitness is indeed constant over time, decreasing ^7^, or increasing ^22^ remains to be elucidated in future studies.

Our mathematical framework, which integrates clonal structure, fitness, and timing through the MACS120 metric, provides a superior prediction of clinical outcomes compared to traditional measurements. This improved predictive power stems from capturing the complete growth trajectory of a clone rather than its current state alone. Our framework’s ability to model clonal cooperation has revealed how mutational context influences fitness effects, exemplified by the differential impact of ASXL1 mutations depending on co-occurring variants. These findings suggest that the binary classification of CH as “benign” or “pre-malignant” oversimplifies a continuous spectrum of risk determined by the confluence of multiple factors.

The implications of this work extend beyond risk stratification. By providing tools for quantifying clonal expansion at the individual level with co-mutational context, our approach enables the investigation of fundamental questions about stem cell biology and aging. The observation that mutation timing correlates with fitness effects suggests age-related changes in selection pressures within the bone marrow microenvironment - a finding that may have broader implications for understanding tissue aging and cancer evolution ^22^.

While our findings represent a significant advance in our understanding of clonal expansion, they also highlight the critical need for more longitudinal studies across diverse populations ^23^. While valuable for identifying associations, cross-sectional analyses cannot capture the dynamic nature of clonal growth or account for individual variation in fitness effects. Future studies should investigate how environmental factors, therapeutic interventions, epigenetics, and genetic background modify clonal dynamics. Additionally, understanding how clone fitness relates to specific molecular mechanisms of expansion may reveal new therapeutic opportunities for preventing progression to malignancy.

This work establishes a framework for transitioning from descriptive to predictive analysis in CH research, marking an important step toward personalized risk assessment and intervention strategies. As we continue to unravel the complexity of clonal evolution, the integration of longitudinal data with sophisticated modeling approaches will be crucial for translating our understanding of CH into improved patient outcomes.

## Materials and Methods

### Participant Samples and Ethics

#### Lothian Birth Cohorts of 1921 and 1936

This study has been approved by NHS Lothian, formerly Lothian Health Board. All participants gave informed consent. Ethics permission for LBC1936 was obtained from the Multicenter Research Ethics Committee for Scotland (wave 1: MREC/01/0/56), the Lothian Research Ethics Committee (wave 1: LREC/2003/2/29) and the Scotland A Research Ethics Committee (waves 2, 3, 4 and 5: 07/MRE00/58). Ethics permission for LBC1921 was obtained from the Lothian Research Ethics Committee (wave 1: LREC/1998/4/183; wave 2: LREC/2003/7/23; wave 3: 1702/98/4/183) and the Scotland A Research Ethics Committee (waves 4 and 5: 10/MRE00/87).

The Lothian Birth Cohorts of 1921 (LBC1921) and 1936 (LBC1936) are two parallel longitudinal studies of cognitive aging recruited from the Lothian area of Scotland, UK. The studies comprise a total of 550 (234:316 male:female; 79.1 median age) and 1,091 (548:543 male:female; 69.5 years median age) participants at baseline in both the LBC1921 and LBC1936, respectively ^9,10^.

Previously, we had identified 73 participants with CH through somatic variant calling of whole-genome DNA sequencing at wave 1 ^24^. In a follow-up study, we used error-sequenced blood from these participants, including a further sixteen where CH had previously been undetected using a targeted gene panel of 75 known myeloid driver mutations across 3-5 time points ^6^. In all, we sequenced 248 samples across 85 participants post-quality control. In this study, we extend our longitudinal cohort by sequencing another 63 participants with previously undetected CH across 3 waves (approximately capturing age at years 70, 76, and 82) in the LBC1936. In all, we have sequenced 189 samples across three batches with six ‘Genome in a Bottle’ (GIAB) controls (two per sequencing batch) allowing us to characterize the background error rates in our sequencing strategy.

#### Women’s Health Initiative (WHI)

The Women’s Health Institute is a prospective study across multiple centers in the US to investigate common disease risk factors (osteoporotic fracturing, cardiovascular disease, etc.) in a cohort of postmenopausal women. In 1993, WHI clinical centers selected women aged between 50 and 79 across the United States who were invited to complete a questionnaire alongside baseline clinical screening, including blood collection ^12^.

A subset of the cohort has been prospectively followed for the preceding quarter century. Additional blood collection occurred at annual visits (AV) one, three, six, and nine-years post-enrollment (AV1, AV3, AV6, AV9). The subsequent WHI Long Life Study (WHI-LLS) facilitated a final visit in 2012-2013, capturing between 14-19 years of late-life (mean 15.4 years). Genomic DNA was sampled from peripheral blood leukocytes at each visit using a 5’ DNA extraction kit.

The longitudinal cohort sampled from the WHI consists of 182 participants without prior history of hematological malignancy and were selected due to, either: (1) having previous whole-genome or target-based sequencing that resulted in the detection of a known CH variant (≥ 2% VAF) in the NHLBI TOPMed project (n = 100), or; (2) having previously collected DNA at three or more waves (n = 86). The authors then use a smMIPS-based assay to target common myeloid driver mutations across the time course (62 years baseline median age; 81 years median age at WHI-LLS) ^8^.

Data for the WHI is available through dbGAP (phs000200.v12.p3). This study underwent review by the WHI Practice and Procedure Committee (project identifier: MS4699P). The authors received epidemiological and variant data upon completing relevant data access requests.

#### Sardinian Longitudinal Project (SardiNIA)

The SardiNIA Longitudinal Project is a study on genetic and epidemiological factors for age-related traits and diseases in the Sardinian population, a homogeneous founder population in Southern Europe. The SardiNIA cohort comprises about 8,000 general population participants (57% females, 43% males), ranging from 18 to 102 years, native of the central east coast of Sardinia, Italy. The study was initiated in 2001, and all cohort members are deeply genetically characterized and granularly phenotypically profiled. Participants signed informed consent to study protocols approved by the Sardinian Regional Ethics Committee (protocol no. 2171/CE).

The SardiNIA cohort captured 5 waves of epidemiological and blood sample data collected over 20 years. The original authors analyzed 385 individuals where serialized data was most readily available. Targeted enrichment of whole blood was then performed with a custom probe set (Agilent SureSelect ELID 3156971) that encompasses 56 known CH and myeloid driver mutations before sequencing ^7,11^.

Data from this study is available through dbGAP (phs000313.v3.p2). Ethical permission for the original study was granted by The East of England (Essex) Research Ethics Committee (REC reference 15/EE/0327).

This study complies with all relevant ethical regulations.

#### Targeted, error-corrected sequencing and data filtering

Our expanded cohort longitudinally sampled from the LBC1936 was processed as per our previous work. Briefly, DNA was extracted from whole blood with Ethylenediaminetetraacetic acid (EDTA) using the Nucleon BACC3 kit (Sigma-Aldrich, GERPN8512), according to the manufacturer’s protocol. Libraries were generated from 200 ng of each DNA sample using the Invitea/Archer VariantPlex® 75 Myeloid gene panel and the VariantPlex® Somatic Protocol for Illumina sequencing (Invitae, AB0108, and the VariantPlex®-HCG Myeloid Kit; see Supplementary Table 2), with adaptations for detecting low variant allele frequencies. Sequencing was performed on the NextSeq 2000 platform (Illumina) using the NextSeq 1000/2000 P3 reagents (300 cycles) v3 kit. One ‘Genome in a Bottle’ (GIAB) DNA sample (NA12878, Coriell Institute) was included in each sequencing batch to ensure reproducibility and support background error modeling and batch correction.

Reads were quality-filtered with a ‘phred’ score threshold of ≥30, and adapter sequences were removed using Trimmomatic (version 0.27) ^25^. Afterward, guided alignment to the human genome assembly hg19 was performed using both bwa-mem (version 0.7.17) and bowtie2 (version 2.2.1) ^26,27^. Unique molecular barcodes, introduced before PCR amplification, were utilized for deduplication to enable accurate multiplexed quantification and reliable mutation detection. Variant calling within targeted regions was conducted using three different tools: Lofreq (version 2.1.0) ^28^, Freebayes ^29^, and Vision (ArcherDX version 6.2.7, unpublished). A consensus was generated by combining the results and selecting for agreement from all three variant callers (see Supplementary Table 3).

All filtered variants with a variant allele frequency (VAF) of 1% or 2% satisfied the following conditions: (1) the number of reads supporting the alternative allele exceeded the minimum coverage requirement without displaying directional bias (AO ≥ 5, UAO ≥ 3); (2) the variants were significantly underrepresented in the Genome Aggregation Database (gnomAD; P ≤ 0.05); (3) the variants were unlikely to be germline, as indicated by a stable VAF (50% or 100%) across all sequencing waves, which could have been underrepresented in gnomAD due to the limited geographic diversity of the LBC cohort; and (4) the variants included events, such as frameshift duplications and deletions, that were overrepresented in the dataset, likely reflecting misaligned artifacts from the capture method (see Supplementary Table 3) ^30^. Additionally, the list was manually curated, cross-referencing variants previously reported in COSMIC, Jaiswal et al., or other published studies ^31^. Finally, for any variant that met the above criteria with a VAF of ≥ 1% or 2% at any time point, all participant-matched data points were included, regardless of baseline VAF.

We received files describing the longitudinally measured variant calls from two existing published longitudinal studies of CH - the SardiNIA and WHI ^7,8^. Despite differences in the sequencing protocols, capture methods, and variant processing steps employed, we found that the enrichment and trajectories of gene mutations were highly concordant across the cohorts. While each of the sequencing methods employed in each cohort targets different gene panels, we estimate that the overlap across all target panels will capture over 90% of the population prevalence of CH in commonly known driver mutations (as estimated from five large cross-sectional studies; Supplemental Table 2) ^4,5,19,32,33^.

#### Mathematical model of clonal dynamics to infer fitness and clinical metrics

We used the mathematical framework we developed to model clonal dynamics and infer fitness from longitudinal variant allele frequency (VAF) data ^6^. Briefly, this framework uses a birth-death process to model stem cell dynamics, where mutations can confer a fitness advantage, *s*, leading to an exponential growth of mutant clones. For each participant with multiple mutations, we considered alternative models representing different possible clonal structures - from all mutations occurring independently to various combinations of co-occurring mutations (see Supp. Methods). Each model defines a specific arrangement of mutations into clones, with mutations in the same clone sharing identical fitness effects and growth dynamics. We then computed the probability of each model conditional on the data to identify the most likely clonal structures and reported the posterior distributions of fitness for this model.

Further, for each individual, we used the preferred clonal structure and the maximum a posteriori (MAP) of each clone’s fitness to predict the age of mutation acquisition (ATMA) using a deterministic model of clonal growth (see Supp. Methods). More specifically, ATMA was determined by maximizing the likelihood of the trajectory given the inferred fitness and a fixed population of wild-type stem cells of (N=100,000), resulting in the most probable time in the past when the clone size was equal to a single cell.

To develop a clinically relevant metric that combines both fitness and timing information, we introduced MACS 120 (Maximum Clone Size at age 120) - the predicted clone size a mutation would reach by age 120. For a mutation with fitness *s* acquired at time ATMA, MACS 120 is calculated as:

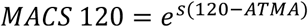

This metric provides a single value that captures both the growth potential of a mutation (through *s*) and the time available for expansion (through ATMA). Further, both fitness and ATMA are conditional on the optimal clonal structure, thereby offering a more comprehensive assessment of a clone’s clinical impact than each parameter alone (Supplemental Table 4).

#### Differential VAF evolution based on mutational context

Recall that our inference model accounts for indirect clonal interactions through the denominator of the variant allele frequency (VAF) calculation. That is, the time evolution of the VAF of variant *j* is deterministically modelled

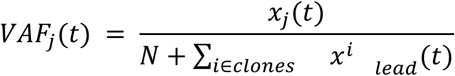

where 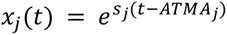 represents the clone size at time *t*, *N* represents the total population of wild-type cells (fixed at 100,000). Further, the sum in the denominator runs over all independent clonal structures in an individual, *i*, and 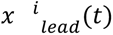 represents the size of the leading mutation

(highest VAF) in the structure.

To quantify the impact of clonal composition within individuals on the VAF growth trajectories, we compared this composition-aware model to an unconstrained model where each mutation grows independently:

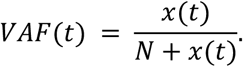

We evaluated the magnitude of clonal competition by predicting VAF levels at age 120 using both the deterministic models above and calculating the difference in predicted VAF for each mutation. This approach allowed us to directly quantify how an individual’s clonal composition affects the perceived growth trajectory of each mutation.

#### Definition of gene categories

Mutated genes were categorized into five functional groups based on their primary molecular functions:

1. Epigenetic Regulation (ER): Genes involved in DNA methylation (DNMT3A, TET2), histone modification (ASXL1, EZH2, CREBBP, KDM6A, PHF6), and other epigenetic processes (IDH1, IDH2).
2. RNA Splicing and Processing (RS): Core splicing factors (SF3B1, SRSF2), splice site recognition factors (U2AF1, U2AF2), and other splicing regulatory proteins (ZRSR2, LUC7L2, PRPF40B, SF3A1).
3. Signal Transduction (ST): Protein kinases (JAK2, BRAF), GTPases (KRAS, NRAS), phosphatases (PPM1D, PTPN11), and other signaling components (CBL, NOTCH1, GNAS, GNB1, SH2B3, NF1, MPL, PTEN, CALR).
4. Transcription Factors (TF): Sequence-specific DNA binding proteins (TP53, BCORL1, WT1, ZNF318, CUX1, CEBPA, RUNX1, ZBTB33, ETV6, GATA2).
5. Chromatin Structure (CS): Genes involved in chromatin organization and chromosome cohesion (BRCC3, STAG2, RAD21, SMC1A, SMC3, CTCF).

This classification system enabled systematic analysis of mutation effects across related biological pathways and facilitated the investigation of pathway-specific fitness effects and co-mutation patterns. Gene assignments were based on established functional annotations from literature and gene ontology databases.

#### Clonal network analysis

We constructed network visualizations to analyze patterns of co-occurrence between mutations. Nodes represent individual mutations, with node size proportional to mutation frequency and node color indicating mutation log fitness (scaled between minimum and maximum log fitness values using a plasma colormap). Edges connect co-occurring mutations, with edge width proportional to co-occurrence frequency and edge color representing the average log fitness of connected mutations. Network visualization was implemented using the ‘pyvis’ library with a force-directed layout. Statistical significance of fitness differences between isolated and co-occurring mutations was assessed for log fitness using the Brunner-Munzel test with Benjamini-Yekutieli correction for multiple comparisons.

#### Survival Analysis

Survival analysis was performed using Cox proportional hazards models implemented through the lifelines Python package. The models incorporated, Maximum VAF (z-score normalized), Maximum fitness (z-score normalized), Age at first observation (z-score normalized), Sex, Maximum predicted clone size at age 120 (MACS 120, z-score normalized). Model performance was evaluated using concordance index and log-rank tests. Age stratification was performed using bins of 50-59, 60-69, 70-79, and 80+ years. Cohort-specific analyses controlled for cohort-specific covariates while maintaining the core model structure.

#### Longitudinal analysis of Blood Markers

We first designed a panel of blood markers (n=43, see Supplementary Table 5) based on previously reported associations in CH and available data across cohorts ^23^. Then we standardized blood marker measurements to enable comparison across different markers and cohorts. For each marker, we first identified and removed outliers if they exceeded a threshold of 3 times the interquartile range. Then we selected the optimal transformation (log, square root, Box-Cox, or Yeo-Johnson) so that the transformed data would be normally distributed within each cohort. Transformed values were converted to z-scores by subtracting the cohort mean and dividing by the cohort standard deviation. Binary markers (those with only two distinct values) were excluded from this transformation process (Supplemental Table 5).

Following standardization, trajectories were analyzed using linear mixed models implemented through ‘statsmodels’, with participant ID as a random effect. For each marker, the model form was:

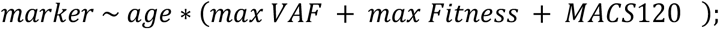

with parameters Maximum VAF, Maximum fitness, and MACS120 were z-score normalized (Supplemental Table 5).

## Data Availability

We have deposited all data, including de-identified FASTQ reads and processed variant calls into NCBI Geo under accession identifier GSE288742 (Reviewer Access Token: elafyywitbwvbqv). Prior targeted longitudinal CH sequencing in the LBCs from our 2022 study is available at NCBI Geo: GSE178936. LBC phenotypic data can be accessed through the Database of Genotypes and Phenotypes (dbGAP) under accession number phs000821.v1.p1. Additional Lothian Birth Cohort data are available in dbGAP or through the LBC Data Access Collaboration (https://www.ed.ac.uk/lothian-birth-cohorts/data-access-collaboration). This platform provides comprehensive information about the cohort, including its background, data summary tables for both LBC1921 and LBC1936, with forms and contact details for data access requests. For inquiries, contact Simon Cox (https://www.ed.ac.uk/profile/simon-cox, simon <DOT> cox <AT> ed <DOT> ac <DOT> uk); a response is typically provided within one month.

Additional secondary datasets included in this analysis can be found at NCBI dbGAP at identifiers phs000313.v3.p2 and phs000200.v12.p3 for the SardiNIA and WHI cohorts. Targeted sequencing data for the SardiNIA cohort is available at the European Phenome-Genome Archive (EGA) with accession numbers EGAD00001007683 and EGAD00001007682.

## Code Availability

All code is available at the following GitHub repository: https://github.com/elc08/Clonal-Haematopoiesis. This contains a detailed schematic describing the dependencies required to run our method, and a schematic that describes the workflow.

## Acknowledgement

We thank all Lothian Birth Cohort (LBC) study participants and research team members who have contributed, and continue to contribute, to ongoing studies. We also would like to thank all cohort participants in the WHI and SardiNIA projects as well as study team members for their previous and ongoing contributions. Finally, we would also like to thank Paul Redmond (University of Edinburgh) for helping us provision data from the LBC and Peter McHardy (CRUK Scotland Institute) for his IT skills and general advice.

## Funding Sources and Conflicts of Interest

K.K. was funded by Blood Cancer UK (grant reference 23001), by an EHA Bilateral grant number ID: BCG-202209-02649 and by the Mayo Clinic Robert and Arlene Kogod Center on Aging and the Mayo Clinic Department of Hematology.

E.L.C. was partially supported by funding from the University of Edinburgh and the Medical Research Council (MC_UU_00009/2) and by K.K EHA Bilateral grant.

E.L.C has been funded by the Spanish Research Agency (AEI), through the Severo Ochoa and Maria de Maeztu Program for Centers and Units of Excellence in R&D (CEX2020-001084-M).

E.L.C thanks CERCA Program/Generalitat de Catalunya for institutional support.

T.C. was funded by the Mayo Clinic Robert and Arlene Kogod Center on Aging.

E.G.K. was funded by the Turkish Ministry of National Education (MEB), YLSY program.

S.E.H is supported by a National Institutes of Health (NIH) research grant (U01AG083829).

S.R.C is supported by a Sir Henry Dale Fellowship jointly funded by the Wellcome Trust and the Royal Society (221890/Z/20/Z).

M.C.H received consulting fees from Comanche Biopharma (modest), research funding from Genentech, site principal investigator for Novartis.

The LBC1921 was supported by the UK’s Biotechnology and Biological Sciences Research Council (BBSRC), The Royal Society, and The Chief Scientist Office of the Scottish Government. LBC1936 is supported by the BBSRC, and the Economic and Social Research Council [BB/W008793/1], Age UK (Disconnected Mind project), the Milton Damerel Trust, and the University of Edinburgh.

---

**Figure S1:**
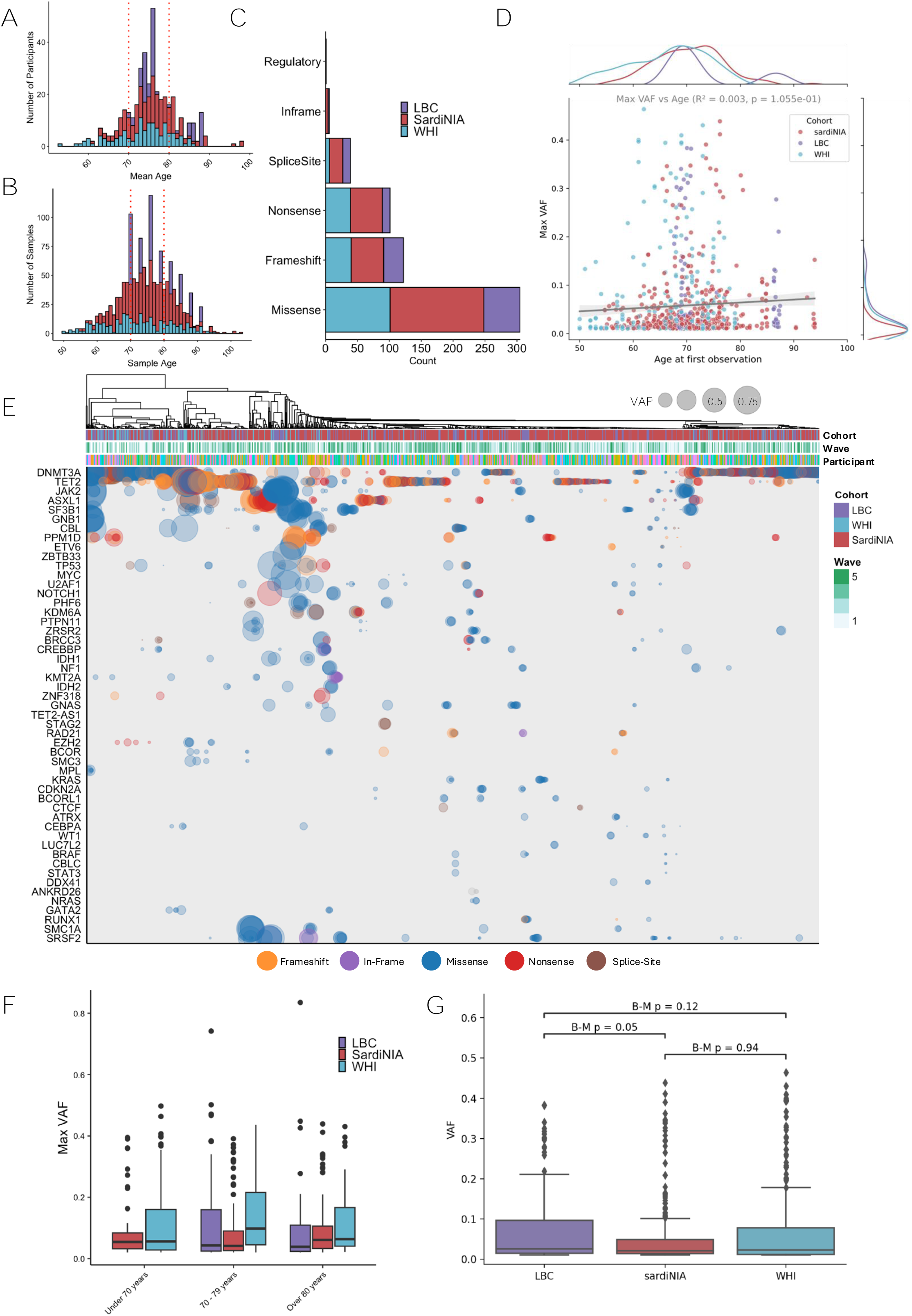
Complete cohort composition and overview. **A.** Mean age of participants (across all measured longitudinal time points), colored by cohort. **B.** Age of participants at sampling points, colored by cohort. **C.** Counts of the functional consequences of variants, colored by cohort. **D.** Age at the first observation of a given variant vs the maximum VAF measured across its trajectory, colored by cohort and flanked by cohort-normalized KDE curves on each axis. We observe a non-significant increase in maximum VAF (Max VAF) with age (R^2^=0.03; p=1.055 e-01). **E.** Variant allele fractions in all mutated genes, showing samples from all participants across all time points clustered by mutated gene. Here, each dot indicates the presence of the largest mutation within a gene for a given individual, which is scaled by VAF and colored by predicted functional consequence. The annotation bar indicates the participant, cohort, and wave number of samples. **F.** VAF levels across all individuals at the first observed time point. Points are colored by cohort, with log_10_ scaled y-axis. **G.** Distribution of maximum VAF (Max VAF) in each participant by cohort and age group. Data is split across age groupings that capture the under 70, 70-79 and 80 years and above. Boxes show the median and exclusive interquartile range with dots showing the outliers. **H.** Distribution of maximum VAF (Max VAF) in each participant by cohort. Boxes show the median and exclusive interquartile range with dots showing the outliers. Statistical significance of VAF differences was assessed using the Brunner-Munzel test.

**Figure S2:**
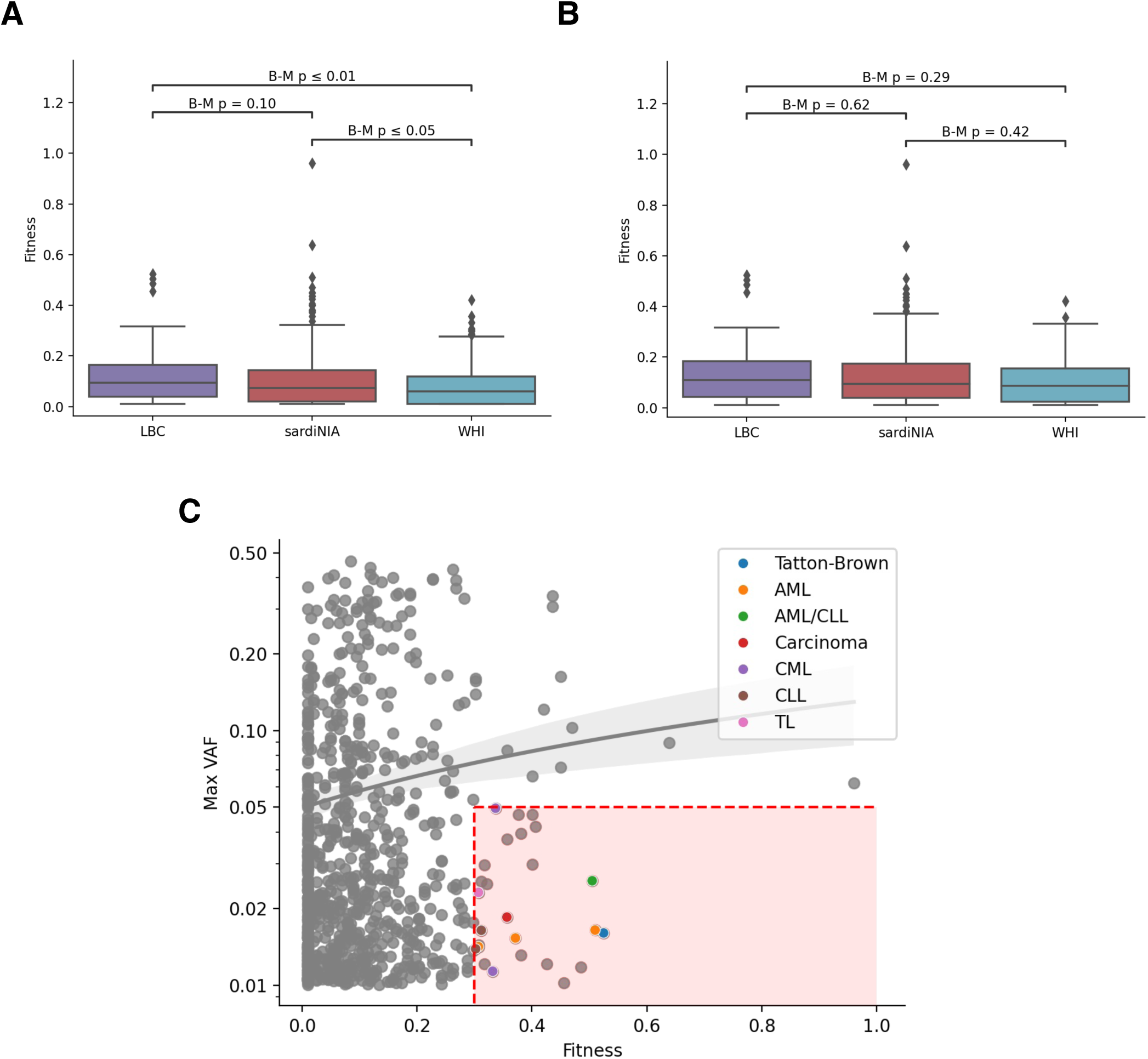
Cohort-specific analyses and mutation characteristics. **A.** Fitness for all mutations in each cohort. Boxes are colored by cohort and show exclusive interquartile range, with diamonds showing outliers. Statistical significance of fitness differences between cohorts was assessed using the Brunner-Munzel test. **B.** Distribution of maximum fitness observed in each participant by cohort. Boxes are colored by cohort and show exclusive interquartile range, with diamonds showing outliers. Statistical significance of fitness differences between cohorts was assessed using the Brunner-Munzel test. **C.** Maximum observed VAF of each mutational trajectory (shown on a log-scale) versus fitness. Colored points indicate variants reported for disease associations, with color showing which disease as shown in legend. Solid grey line and shaded region show linear regression fit showing a positive correlation between max observed VAF and fitness (R=0.1, p=0.002). Y-axis is log transformed to increase the visibility of low VAF variants. AML: Acute Myeloid Leukemia; CML: Chronic Myeloid Leukemia; CLL: Chronic Lymphoid Leukemia; TL: T-cell Lymphoma.

**Figure S3:**
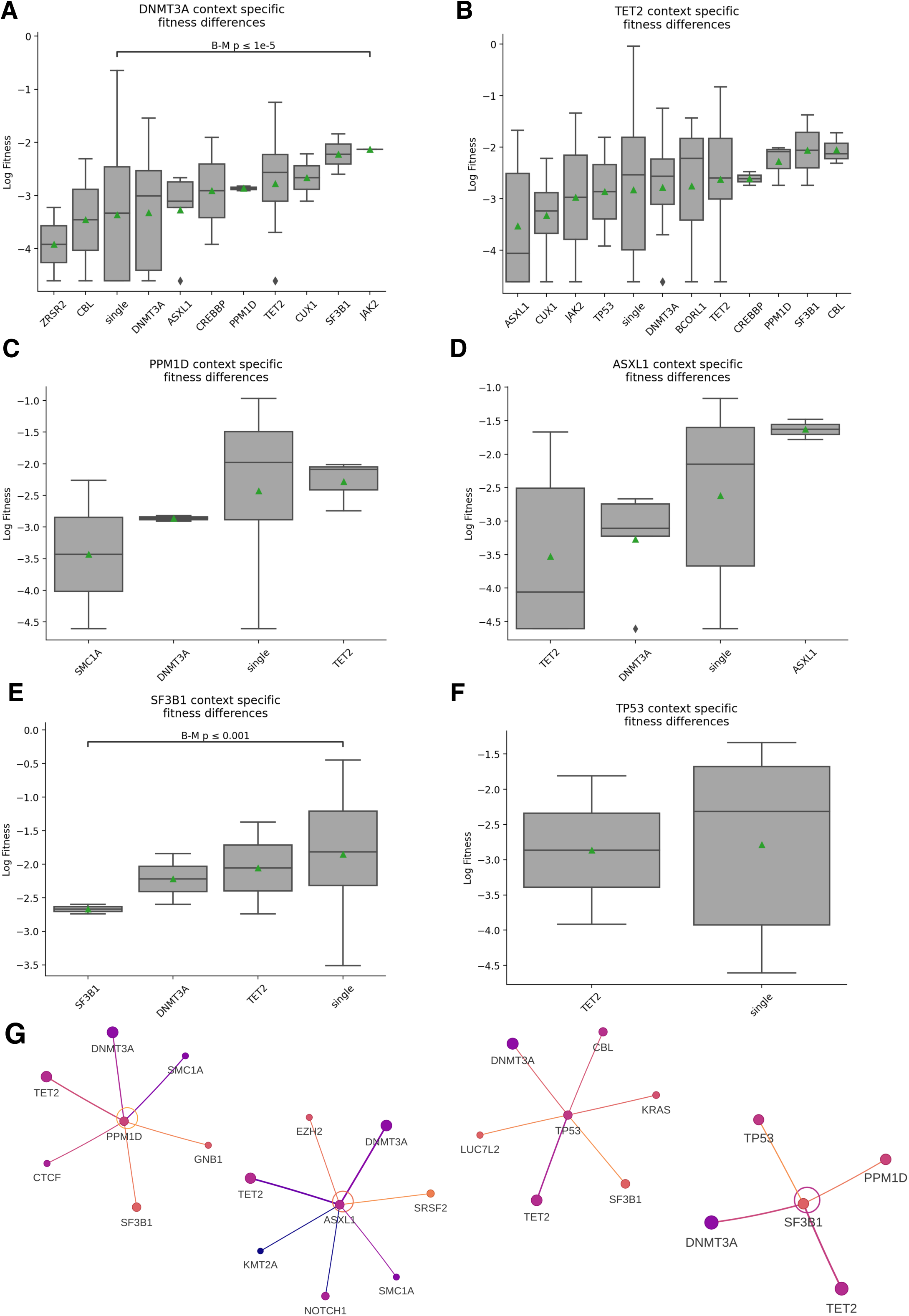
Differential fitness for different clonal compositions. **A-F.** Differential fitness analysis of mutations in a reference gene (**A.** DNMT3A, **B.** TET2, **C**. PPM1D, **D.** ASXL1, **E.** SF3B1, **F.** TP53) across different combinations of co-mutated genes. For each reference gene we show boxplots comparing the log fitness of singly occurring mutations in a clone and co-occurring with mutations in other genes (only shown if more than 1 instance is observed). Boxes show median and exclusive interquartile range and diamonds show outlier values. Statistical significance of fitness differences between isolated and co-occurring mutations was assessed for log fitness using the Brunner-Munzel test with Benjamini-Yekutieli correction for multiple comparisons. **G.** Network visualization showing mutation co-occurrence patterns for a single central gene. This representation is a direct isolation of the network associated to a single node as displayed in Fig 3C. Nodes represent individual mutations, with node size proportional to log counts of mutation instances and node color indicating average mutation log fitness (scaled between minimum and maximum log fitness values). Edges connect co-occurring mutations in a clonal structure, with edge width proportional to co-occurrence log counts and edge color representing the average log fitness of connected mutations.

**Figure S4:**
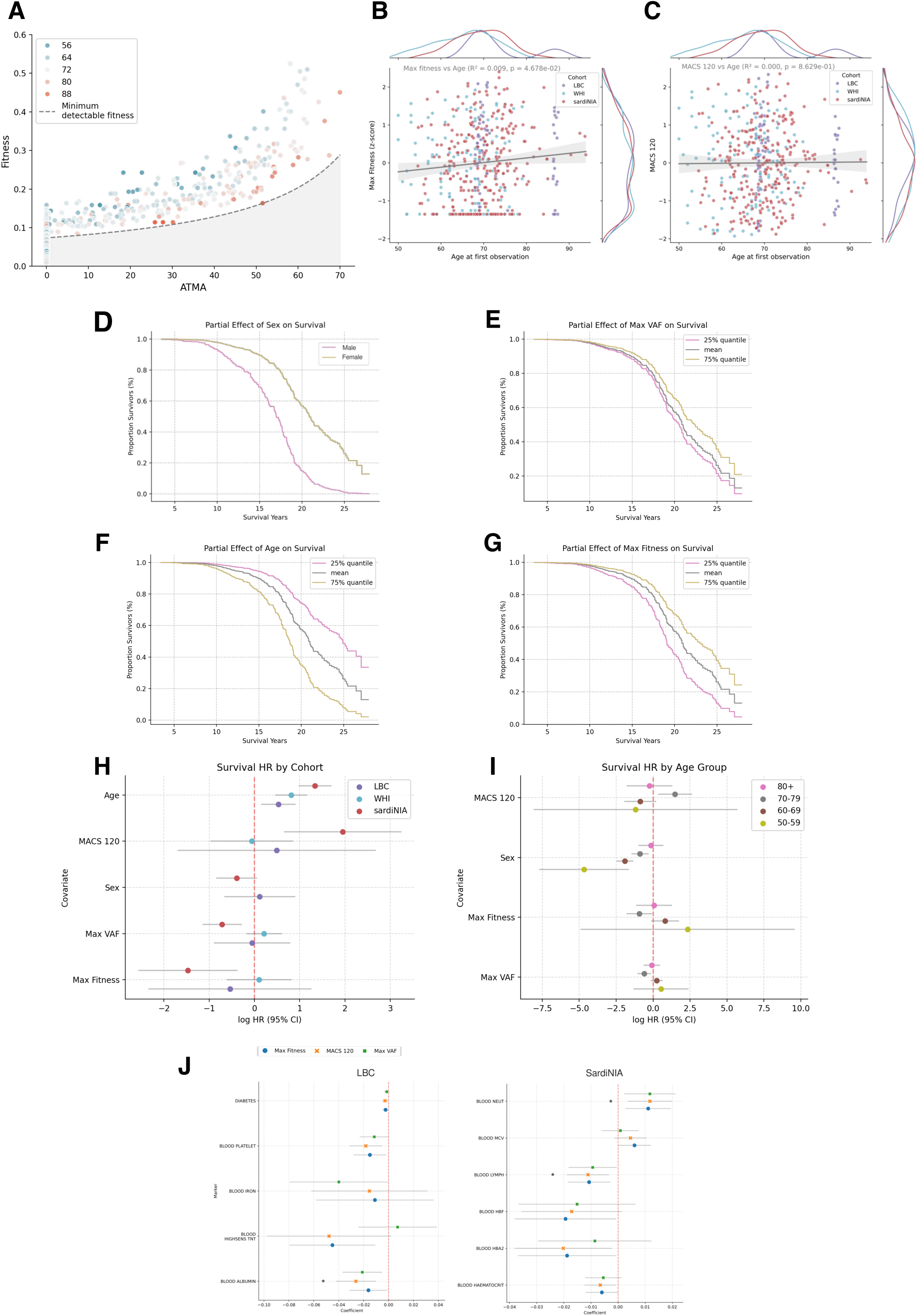
MACS120 and clinical outcomes. **A.** Maximum fitness versus Age at Time of Mutation Acquisition (ATMA) with points colored by age of participant at first observation. The absence of low-fitness high-ATMA variants is due to detection limitations (dashed line, see Methods). **B.** Correlation between maximum observed fitness in each individual and their age at first observation, with points colored by cohort. Grey line and shaded area show linear fit (R^2^ =0.009, p=0.04). Marginal distributions shown as cohort-normalized Kernel Density Estimation (KDE) curves on each axis. **C.** Correlation between MACS120 in each individual and their age at first observation, with points colored by cohort. Grey line and shaded area show linear fit (R^2^ =0, p=0.86). Marginal distributions shown as cohort-normalized Kernel Density Estimation (KDE) curves on each axis. **D-G** Partial effects in survival probability over time of model parameters included in the survival analysis. Parameters analyzed are: **D.** Sex, **E.** Maximum observed VAF in participant (z-score normalized) **F.** Age of participant at first observation (z-score normalized) **G.** Maximum observed fitness in participant (z-score normalized)). For each parameter, we show how variations in their values (25% quantile, baseline and 75% quantile) influence survival trajectories while controlling for other variables. **J.** Cohort-specific analysis of the association between Maximum Fitness (blue dot), Maximum VAF (green square) and MACS120 (orange cross) and longitudinal changes in blood markers at the individual level. For each blood marker only metrics that showed a significant correlation are shown (see Methods). We display effect size and confidence intervals with a grey line. Acronyms: Lymphocytes (LYMPH) High-Sensitivity Cardiac Troponin T (HIGHSENS TNT), Mean Corpuscular Volume (MCV), Neutrophils (Neut), Thyroxine (T4).

## References

1. Lee-Six, H. et al. Population dynamics of normal human blood inferred from somatic mutations. Nature 561, 473–478 (2018).

2. Florez, M. A. et al. Clonal hematopoiesis: Mutation-specific adaptation to environmental change. Cell Stem Cell 29, 882–904 (2022).

3. Watson, C. J. et al. The evolutionary dynamics and fitness landscape of clonal hematopoiesis. Science 367, 1449–1454 (2020).

4. Jaiswal, S. et al. Age-related clonal hematopoiesis associated with adverse outcomes. N. Engl. J. Med. 371, 2488–2498 (2014).

5. Genovese, G. et al. Clonal hematopoiesis and blood-cancer risk inferred from blood DNA sequence. N. Engl. J. Med. 371, 2477–2487 (2014).

6. Robertson, N. A. et al. Longitudinal dynamics of clonal hematopoiesis identifies gene-specific fitness effects. Nat. Med. 28, 1439–1446 (2022).

7. Fabre, M. A. et al. The longitudinal dynamics and natural history of clonal haematopoiesis. Nature 606, 335–342 (2022).

8. Uddin, M. M. et al. Longitudinal profiling of clonal hematopoiesis provides insight into clonal dynamics. Immun. Ageing 19, 23 (2022).

9. Deary, I. J. et al. The Lothian Birth Cohort 1936: a study to examine influences on cognitive ageing from age 11 to age 70 and beyond. BMC Geriatr. 7, 28 (2007).

10. Deary, I. J., Gow, A. J., Pattie, A. & Starr, J. M. Cohort profile: the Lothian Birth Cohorts of 1921 and 1936. Int. J. Epidemiol. 41, 1576–1584 (2012).

11. Orrù, V. et al. Genetic variants regulating immune cell levels in health and disease. Cell 155, 242–256 (2013).

12. Design of the women’s health initiative clinical trial and observational study. Control. Clin. Trials 19, 61–109 (1998).

13. 13. Mon Père, N., Terenzi, F. & Werner, B. The dynamic fitness landscape of ageing haematopoiesis through clonal competition. *BioRxiv* (2024) doi:10.1101/2024.04.16.589764.

14. 14. MacGregor, H. A., Easton, D. F. & Blundell, J. R. Modelling the age-related deceleration of clonal haematopoiesis in UK Biobank. *BioRxiv* (2023) doi:10.1101/2023.12.21.572706.

15. Richardson, D. R. et al. Genomic characteristics and prognostic significance of co-mutated ASXL1/SRSF2 acute myeloid leukemia. Am. J. Hematol. 96, 462–470 (2021).

16. Patnaik, M. M. et al. EZH2 mutations in chronic myelomonocytic leukemia cluster with ASXL1 mutations and their co-occurrence is prognostically detrimental. Blood Cancer J. 8, 12 (2018).

17. Bolton, K. L. et al. Cancer therapy shapes the fitness landscape of clonal hematopoiesis. Nat. Genet. 52, 1219–1226 (2020).

18. Arends, C. M. & Jaiswal, S. Gene-Specific Machine Learning Models to Classify Driver Mutations in Clonal Hematopoiesis. Cancer Discov. 14, 1581–1583 (2024).

19. McKerrell, T. et al. Leukemia-associated somatic mutations drive distinct patterns of age-related clonal hemopoiesis. Cell Rep. 10, 1239–1245 (2015).

20. 20. Walkowiak, B., MacGregor, H. A. & Blundell, J. R. No evidence of immunosurveillance in mutation-hotspot driven clonal haematopoiesis. *BioRxiv* (2024) doi:10.1101/2024.09.27.615394.

21. Bowman, R. L., Busque, L. & Levine, R. L. Clonal hematopoiesis and evolution to hematopoietic malignancies. Cell Stem Cell 22, 157–170 (2018).

22. Heyde, A. et al. Increased stem cell proliferation in atherosclerosis accelerates clonal hematopoiesis. Cell 184, 1348–1361.e22 (2021).

23. Kar, S. P. et al. Genome-wide analyses of 200,453 individuals yield new insights into the causes and consequences of clonal hematopoiesis. Nat. Genet. 54, 1155–1166 (2022).

24. Robertson, N. A. et al. Age-related clonal haemopoiesis is associated with increased epigenetic age. Curr. Biol. 29, R786–R787 (2019).

25. Bolger, A. M., Lohse, M. & Usadel, B. Trimmomatic: A flexible trimmer for Illumina sequence data. Bioinformatics 30, 2114–2120 (2014).

26. Li, H. & Durbin, R. Fast and accurate short read alignment with Burrows-Wheeler transform. Bioinformatics 25, 1754–1760 (2009).

27. Langmead, B. & Salzberg, S. L. Fast gapped-read alignment with Bowtie 2. Nat. Methods 9, 357–359 (2012).

28. Wilm, A. et al. LoFreq: a sequence-quality aware, ultra-sensitive variant caller for uncovering cell-population heterogeneity from high-throughput sequencing datasets. Nucleic Acids Res. 40, 11189–11201 (2012).

29. 29. Garrison, E. & Marth, G. Haplotype-based variant detection from short-read sequencing. *arXiv* (2012).

30. Karczewski, K. J. et al. The mutational constraint spectrum quantified from variation in 141,456 humans. Nature 581, 434–443 (2020).

31. Tate, J. G. et al. COSMIC: the catalogue of somatic mutations in cancer. Nucleic Acids Res. 47, D941–D947 (2019).

32. Zink, F. et al. Clonal hematopoiesis, with and without candidate driver mutations, is common in the elderly. Blood 130, 742–752 (2017).

33. Young, A. L., Challen, G. A., Birmann, B. M. & Druley, T. E. Clonal haematopoiesis harbouring AML-associated mutations is ubiquitous in healthy adults. Nat. Commun. 7, 12484 (2016).

